# Analysis the Detailed Process of Glomerular Capillary Formation Using Immunofluorescence Perform With Ultrathick Section

**DOI:** 10.1101/810382

**Authors:** Ting Yu, Chi Liu

## Abstract

**Background:** Glomerular capillary formation is one of the fundamental mysteries in renal developmental biology. However, there are still debates on this issue, and its detailed formation process has not been clarified.

**Results:** To resolve this problem, we performed antibody staining with ultra-thick section on embryonic and postnatal mouse kidneys to detect and analyze the development of glomerular capillaries. We found that blood vessel of the fetal kidneys expanded through proliferation and sprouting. During the comma-stage and S-shaped stages, 3-4 capillaries began to bud and migrate into the glomerular cleft, forming a capillary bed in the Bowman’s capsule. Then, the capillary bed expanded into mature glomerular capillary by intussusceptive angiogenesis. The afferent and efferent arterioles were formed through pruning. The distribution of VEGFA in the nephron epithelial cells but not only in podocytes, induced multiple capillaries sprouted into the glomerular cleft. And CXCR4 played an important role in the differentiation and expansion of capillary bed into glomerular capillary.

**Conclusions:** Immunofluorescence performed with ultra-thick section allowed us to investigate the development of complex structure tissues systematically and comprehensively.

## Background

The glomerulus is a highly differentiated microvascular bed. Glomerular afferent arterioles enter the Bowman’s capsule from the vascular pole and are divided into 5 to 8 branches, which are then divided into many looped capillaries. These capillaries are coiled into 5 to 8 capillary lobules or segments, with few anastomotic branches between capillaries. The capillaries of each lobule concentrate in turn into larger vessels, and then converge with other lobular vessels to form outgoing arterioles, leaving the glomerulus from the vascular pole. Endothelial cells of glomerular capillary are fenestrated and glycocalyx-coated. As a blood filter, glomerular capillary allows small molecules and proteins to pass through, while retaining high molecular weight proteins and cells in the blood circulation [1]. The special structure and function of the glomerular capillary suggest that their formation process also has its particularity. However, the precise process of glomerular capillary formation has not been elucidated.

Blood vessels arise by vasculogenesis or angiogenesis. Vasculogenesis indicates that new blood vessels are formed by endothelial progenitor cells (EPCs), whereas angiogenesis is the process of budding new capillaries from preexisting vessels [2]. Whether glomerular capillary is formed by vasculogenesis or angiogenesis is still uncertain. In 1965, Potter E. L. believed that glomerular capillaries originate when a single capillary loop grows into the glomerular cleft in the S-shaped stage. Then, the capillary loop becomes divided into six to eight loops and eventually forming mature glomerular capillaries [3, 4]. However, more recent studies refute this conclusion. When mouse embryonic kidneys are implanted into anterior eye chambers of adult rats, the vast majority of the microvasculature is from the donor. This means that cells from the implanted kidney are responsible for the development of the blood vessels in oculus [5, 6]. Then individual EPCs are identified in the developing kidney using markers such as FLK1. At the S-shaped stage, these EPCs migrate into the glomerular cleft. Once inside the developing glomerulus, EPCs proliferate in situ and aggregate to form the first capillary loops [7, 8]. One of the reasons for these controversies is that the formation of glomerular capillary has not been elucidated.

The previous studies were based on immunohistochemistry and immunofluorescence of approximately 15 μm-thick sections [3, 7]. With these thin sections, we can only obtain a limited portion of the tissue structure. Therefore, we reexamined the development of glomerular capillary by using immunofluorescence performed with ultra-thick section and obtained a detailed process of glomerular capillary development.

## Results

### The origin of renal vascular

The thickness of a glomerulus is approximately 97.4 μm [9]. To record the developmental process of glomerular capillaries more accurately and completely, we developed an immunofluorescence method using ultra-thick section. Sections were generated at 150 μm. Negative control was performed for each antibody used in the following studies, and no non-specific signal was detected (data not shown). Then, we used immunostaining to detect the expression of FLK1 and CD34 which were expessed in EPCs but also in mature blood vessels [7, 10]. Compared with 15 μm tissue sections (Fig. 1a), we found that no individual FLK1 expression (FLK1^+^) or CD34^+^ cells were found in the ultra-thick fetal kidney sections, and all of these endothelial cells were interconnected to form blood vessels (Fig. 1b, c). To determine whether these blood vessels were mature, we detected two mature blood vessels markers, endomucin and CD31 at *E 13* to *E 16* [11, 12]. We observed that CD31 was localized between endothelial cells (Fig. 1d), which demonstrated a stable connection between endothelial cells. Endomucin was widely expressed in renal vessels and increased with the development of embryos (Fig. 1e, f). These data indicate that blood vessels in the embryonic kidney exist in a continuous and mature manner.

**Fig. 1.**
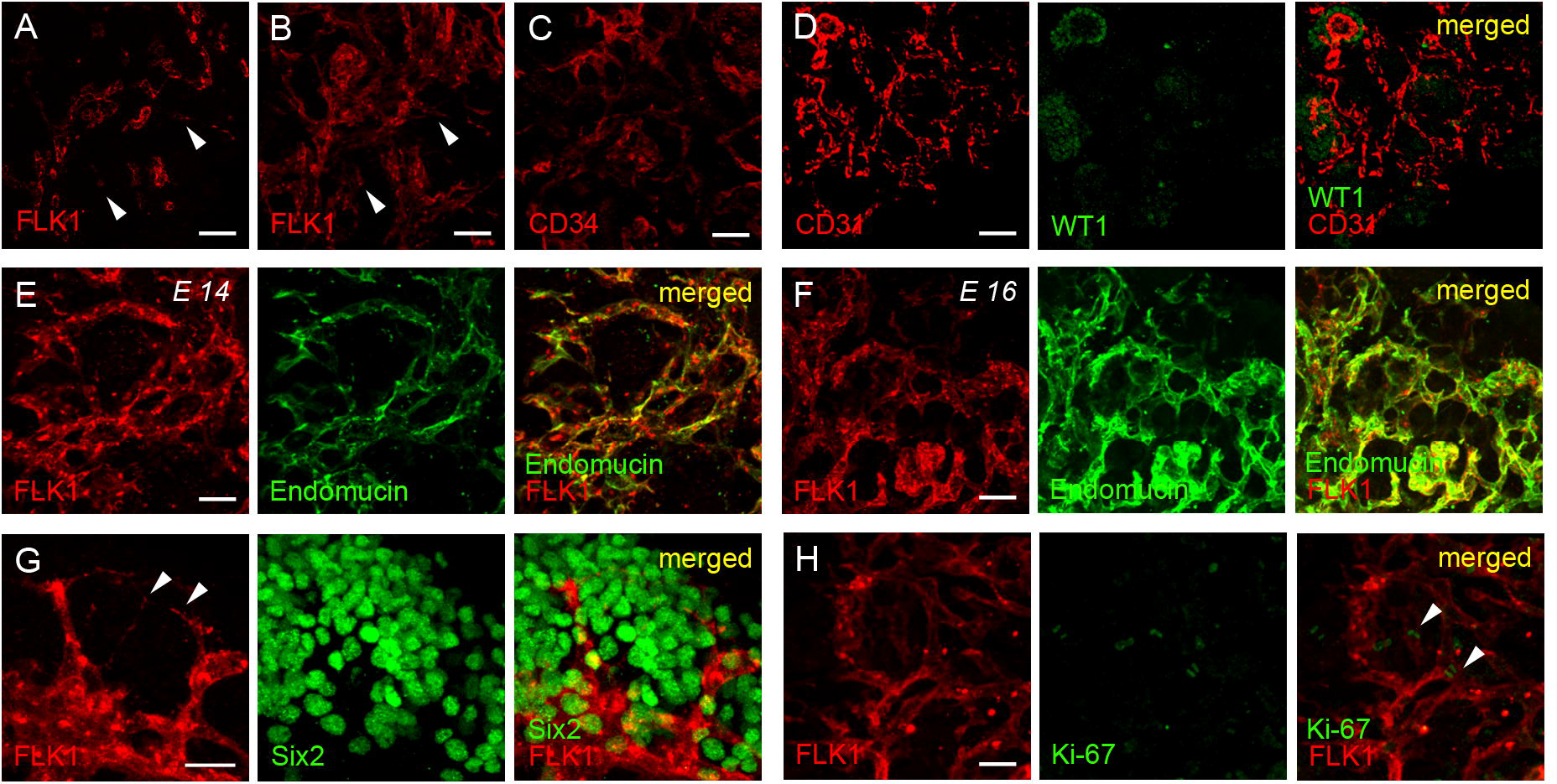
Cellular source of fetal renal vessels. (**a**) FLK1 appears to label individual endothelial cells (arrowheads) when only 15 μm of the 150 μm thick *E 13* kidney sections are displayed. (**b**) In *E 13* kidney sections of 150 μm thickness, FLK1 marks continuous blood vessels. The arrowheads indicate the same locations as A. (**c**) The expression pattern of CD34. CD34 labels continuous blood vessels similar to FLK1. (**d**) The expression pattern of CD31. CD31 staining (left), WT1 staining (middle, expression in podocytes) and the merged image (right) indicate that all blood vessels in the fetal kidney express CD31. (**e**and **f**) The expression pattern of endomucin. FLK1 staining (left), endomucin staining (middle) and the merged image (right) indicate that the expression level of endomucin was relatively low in the *E 14* (**e**) and increased in *E 16* (**f**). (**g**) Six2 staining (middle) indicates the cap mesenchyme, FLK1 staining (left) and the merged image (right) indicate vascular sprouting occurs in the fetal kidney (arrowheads). (**h**) Ki-67 was used to detect the proliferation of renal vessels. FLK1 staining (left), Ki-67 staining (middle) and the merged image (right) reveal that endothelial cells are highly proliferative cells during kidney development (arrowheads). Scale bar = 10 μm.

In that case, what is the source of renal blood vessels in the fetal kidney? Blood vessel sprouting was observed around Six2^+^ cap mesenchyme (Fig. 1g; Additional file 2: Video S1). Sprouting requires endothelial cell proliferation [13]. We detected cell proliferation with Ki-67 [14]. A large number of Ki-67^+^ cells appeared in the renal cortex, of which 27.09% ± 4.32% were vascular endothelial cells (Fig. 1h). This suggests that the source of the vascular endothelial cells in the early stage of kidney development is sprouting and proliferation. These results further confirm a recently published study on renal vascular development [15], which demonstrate that polygonal networks of vessels form in cycles at the periphery of the kidney by angiogenesis. However, the fine development of glomerular capillary need further elucidation.

### Detailed development processes of glomerular capillary

Subsequently, we tracked the development of glomerular capillary with two markers of early glomerulus before and after birth. WT1 can label developing podocytes. NCAM1 is a marker of renal epithelial cells. Nephron begins to appear on the *E 13* and stop producing new nephrons within a few days after birth. During this period, new glomeruli appeared every day. We found that glomerular capillary formed in the same way before and after birth (Additional file 1: Fig. S1, a-i). The nascent nephron epithelium progresses through four stages: renal vesicle, comma-shaped body, S-shaped body, maturation [16]. At first, capillaries formed a network surrounding renal vesicle (Fig. 2a). In the comma and S-shaped stages, 2-4 capillaries budding from the capillary network began to extend into the cleft of the glomerulus (Fig. 2b, c; Additional file 3: Video S2). These capillaries intersected and fused in the cleft, forming a capillary bed with multiple holes (Fig. 2c-f). Sprouting was also found on the capillary bed. The sprouting of these marginal vessels expanded the area of the capillary bed through extension and anastomosis (Fig. 2e, f; Additional file 4 and 5: Videos S3 and S4).

**Fig. 2.**
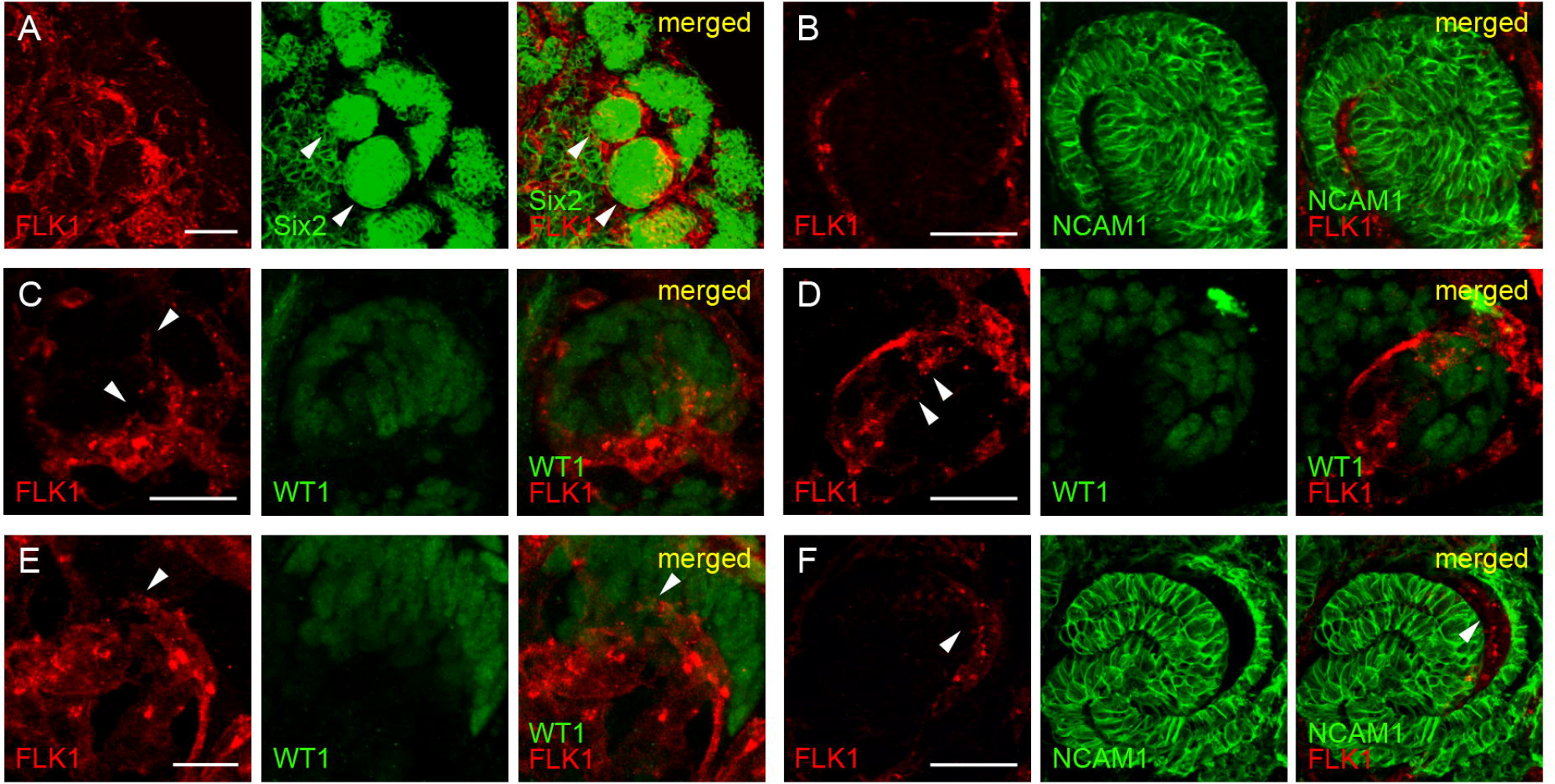
Initial stage of glomerular capillary development. (**a**) The formation of the vesicle capillary network. Ncam1 staining (middle) indicates the renal vesicle (arrowheads), and FLK1 staining (left) and the merged image (right) indicate that capillaries form a capillary network surrounding the renal vesicles. (**b**) The initiation of glomerular capillary development. NCAM1 staining (middle) indicates the renal comma-shaped body. Blood vessel begang extend into the glomerular cleft (arrowheads). (**c**) WT1 staining (middle) indicates Immature podocytes, and FLK1 staining (left) and the merged image (right) indicate that multiple capillaries (arrowheads) extend into the glomerular cleft. (**d**) Capillary bed formation. FLK1 staining (left), and WT1 staining (middle) and the merged image (right) reveal the capillaries (arrowheads) that extend into the glomerular cleft fuse to form a capillary bed. (**e**) The capillary bed enlarges its area by sprouting. The arrowhead indicates the sprouting of blood vessels on the capillary bed. (**f**) Capillary beds were formed in S-shaped body. The arrowhead indicates the capillary bed. Scale bar = 10 μm.

In the process of developing into the Bowman’s capsule, the front end of the S-shaped body extended and encapsulated the capillaries that extend into the glomerular cleft (Fig. 3a). Influenced by the change of glomerular cleft shape, the capillary bed also bended and tended to be bowl shaped (Fig. 3b; Additional file 6: Video S5). Then the capillary bed began to expand and became spherical. During this process, we did not find sprouting from glomerular capillaries (Fig. 3b, c). Previous studies have shown that glomerular capillaries are enlarged via intussusceptive angiogenesis [17, 18]. Intussusceptive angiogenesis is a type of new blood vessel formation in which a capillary is longitudinally split into two vascular channels due to the formation and merging of intraluminal tissue pillars [17]. Ki-67 was used to detect the proliferation of endothelial cells during this period (Fig. 3c). We found that endothelial cells are highly proliferative during the maturation of the glomerulus.

**Fig. 3.**
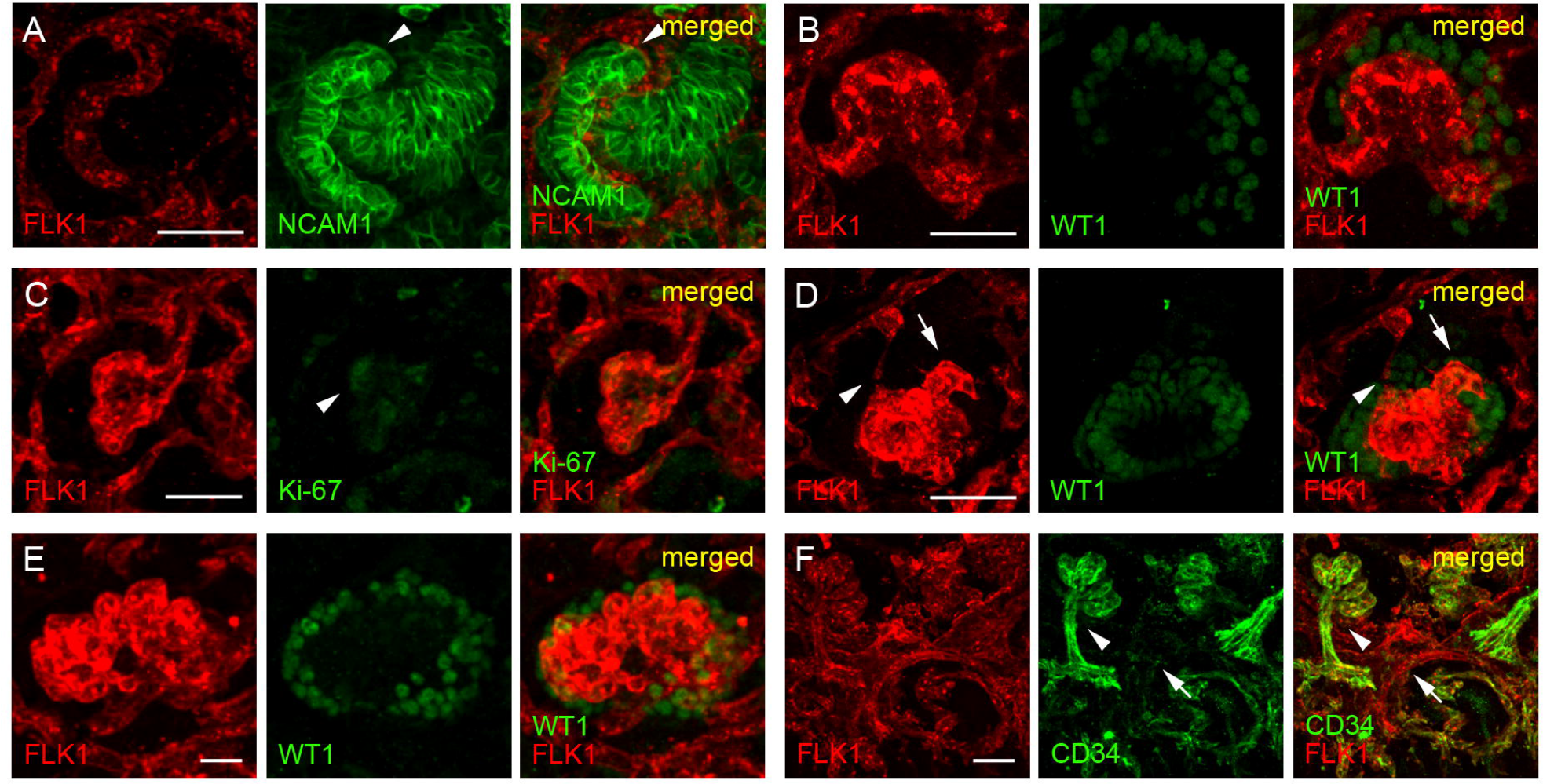
The mature stage of glomerular capillaries. (**a**) The initiation of Bowman’s capsule formation. NCAM1 staining (middle) indicates that the front end of the S-shaped body encapsulates the capillary bed (arrowhead). (**b**) The capillary bed was bent into a bowl shape. (**c**) glomerular capillaries began to expand spherically without sprouting. Ki-67 staining (arrowhead) indicates endothelial cells proliferate rapidly during this process. (**d**) The formation of the afferent and efferent arteriole. Redundant blood vessels (arrowhead) are removed by pruning during the process of glomerular capillary maturation. The remaining blood vessels enlarge and eventually become afferent arteriole or efferent arteriole (arrow). (**e**) The size of the mature glomerular capillaries. The diameter of glomerular capillary on the first day after birth is approximately 70 μm. (**f**) The expression pattern of CD34. The expression of CD34 began to increase gradually in the glomerular artery (arrowhead), while the expression in the peritubular capillaries began to decrease and was lost after glomerular maturation (arrow). Scale bar = 10 μm.

At comma-shaped and S-shaped stages, 2-4 capillaries extended into cleft to form a capillary bed. But the mature glomerulus had only one afferent arteriole and one efferent arteriole [19]. How were these two vessels formed? During the development of the capillary bed toward the mature glomerulus, pruning of superfluous vessels was found [19]. As shown in Figure 3D and Video S6, a blood vessel disconnected and regressed from the glomerulus, while the afferent and efferent arterioles developed into larger vessels. Subsequently, the glomerulus began to further expand, and the largest glomerulus was approximately 70 μm in diameter on the first day after birth (Fig. 3e). During the process of glomerular maturation, the expression of CD34 began to increase gradually in the glomerular artery, while the expression in the peritubular capillaries began to decrease or lost after glomerular maturation (Fig. 3f). Therefore, CD34 can be used as a mature marker of glomerular capillaries.

### Molecular mechanism of glomerular capillary development

The molecular mechanism of glomerular capillary formation is another important research topic. Previous studies have shown that VEGFA is a key factor in development of development of glomerular capillary. Loss of the VEGFA gene from developing podocytes in mice results in reduced glomerular endothelium, but not a complete loss [21, 22]. This suggests that VEGFA has other sources. Using immunofluorescence performed with ultra-thick section, we found that VEGFA was highly expressed in all the epithelial cells at the comma-shaped and S-shaped stages (Fig. 4a, b), This indicates that, in addition to podocytes, VEGFA secreted by other epithelial cells may also responsible for the formation of capillary network around Bowman’s capsules and renal tubules. VEGFA is also the reason for multiple capillaries to enter the glomerular cleft to form the capillary bed (Fig. 4b). CXCR4 is another factor that plays an important role in the development of glomerular capillaries [23]. We found that CXCR4 was mainly expressed in some glomerular endothelial cells during the development of the capillary bed into the mature glomerulus (Fig. 4c, d). Previous studies have shown that *cxcr4* mutants exhibited glomeruli with aneurismal dilation of the capillaries [23]. However, the reason for this structure is not clear. Immunofluorescence performed with ultra-thick section showed that FLK1 decreased significantly in the glomerular vessels of *cxcr4* mutants (Fig. 4e). FLK1 plays a decisive role in the proliferation and differentiation of blood vessels [24]. These results indicate that the glomerular endothelial cells of *cxcr4* mutant did not differentiate correctly in the process of capillary bed expansion into glomerulus.

**Fig. 4.**
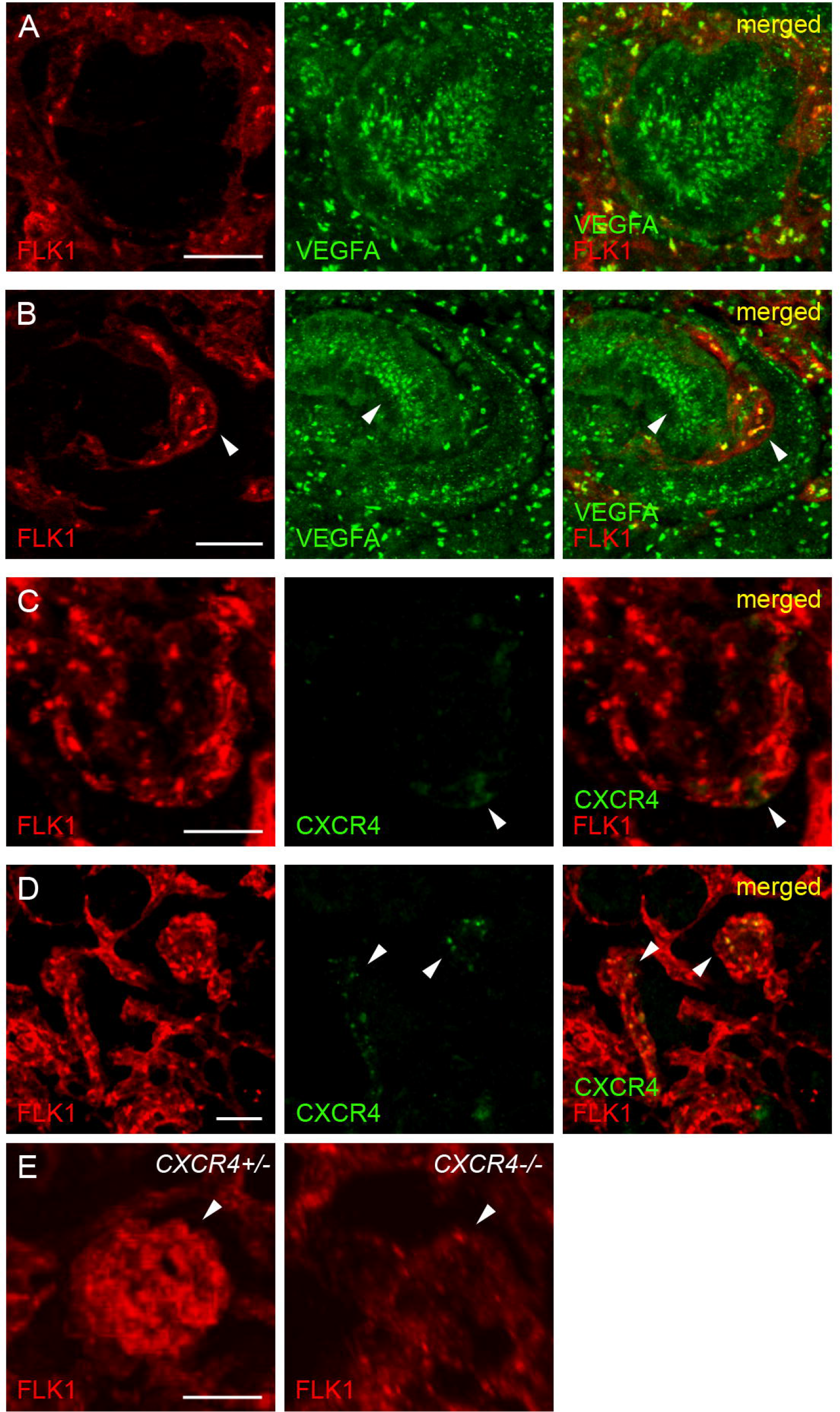
The mechanisms of VEGFA and CXCR4 regulation in glomerular capillary development. VEGFA expression in all epithelial cells of the comma-shaped (**a**) and S-shaped body (**b**) was observed. The expression pattern of CXCR4. CXCR4 was expressed in part of the endothelial cells (arrowhead) during capillary bed (**c**) and glomerular (**d**) expansion. (**e**) Changes of FLK1 expression in the *cxcr4* mutant. The expression of FLK1 in the *cxcr4* mutant glomerular capillaries (right) was significantly decreased by comparing with control (left). Scale bar = 10 μm.

## Discussion

In the traditional methods of studying kidney development, immunohistochemistry and immunofluorescence are usually performed in tissue sections with a thickness of approximately 15 μm [5, 18]. However, the thickness of the nephron is generally more than 100 μm, which means that traditional methods unable obtain the overall structure of the nephron. In addition, because of the low transparency of tissues and the limited penetration depth of antibodies, whole-mount immunofluorescence can only obtain information on the surface of tissues. Tissue optical clearing techniques can be used to obtain immunofluorescence images of deeper tissues, However, this technology requires special instruments and long-term treatment, and the morphology and fluorescence of tissues can not be completely maintained [25]. With the development of imaging technology such as high resolution confocal microscopy, we have been able to obtain three-dimensional images of normal tissues over 100-300 um [26, 27]. Embryonic tissues are more transparent than adult tissues, which enables us to obtain deeper tissue images. Therefore, many tissue structures can be fully acquired in this case. Using ultra-thick section for three-dimensional imaging can help us to obtain as complete information as possible. Therefore, immunofluorescence performed with ultra-thick section is an alternative method for studying the development of tissues with complex structures.

Previous studies used FLK1, CD34 and other proteins as markers of EPCs [10, 28]. Now, with the renewal of interest in vascular development, we know that these genes are expressed not only in EPCs but also in mature blood vessels [29, 30]. Using immunofluorescence performed with ultra-thick section, we observed that endothelial cells are found in a continuous state during the embryonic stage of the kidney, rather than as individual EPCs. The self-proliferation and sprouting of blood vessels provide the source of blood vessels in the fetal kidneys. We demonstrated that VEGFA secreted by renal epithelial cells induced multiple capillaries sprouted into the glomerular cleft and formed a capillary bed, which is the basis of glomerular capillary formation. Then, the capillary bed expanded into mature glomerular capillaries by intussusceptive angiogenesis. CXCR4 determined the differentiation of endothelial cells in this process. Pruning played an important role in the formation of the afferent and efferent arterioles. The mechanisms of pruning excess blood vessels in glomerular development will be a meaningful future research topic.

### Conclusions

Taken together, we have studied the cellular and molecular mechanisms of the development of glomerular capillary by immunofluorescence performed with ultra-thick section and found glomerular capillary formation by angiogenesis. This study may provide clues to the repair of damaged glomerular capillary and the vascularization of tissue engineered kidney.

## Methods

### Animals

All mice used to generate embryos were bred into the C57BL/6 background (purchased from Beijing, China). To generate the cxcr4 mutant, UBC-Cre-ERT2 mice (purchased from Shanghai, China) were crossed with CXCR4 Flox mice (provided by Professor Bi-Sen Ding, Sichuan University); the morning of discovery of the vaginal plug was designated embryonic day 0.5 (*E 0.5)*. To induce *cxcr4* knockout, the pregnant females were injected with 1 mg of tamoxifen and 1 mg of progesterone diluting in 100 μL of corn oil at *E* and *E 11.5*. Animals were housed in a standard animal facility under controlled temperature (22°C) and with free access to water and mouse chow. All animal work has been conducted according to the ethics guidelines of the Army Medical University of China and and the experimental procedures were carried out in accordance with the Guide for the Care and Use of Laboratory Animals published by the National Institutes of Health (NIH Publication No. 85–23, revised 1996). This study was approved by the research ethics committee of the Army Medical University of China.

### Ultra-thick sections

Embryos were allowed to develop in utero up to *E 12* before harvesting tissues. Mice were sacrificed with sodium pentobarbital via an intraperitoneal injection (60 mg/kg). Dissected kidneys were fixed in 4% formaldehyde overnight at 4°C, and then respectively soaked in 15% and 30% sucrose overnight at 4°C. After embedding in OCT (Sakura, ZLI-9302), cryosections were generated at 150 μm using Microm HM 550 cryostat.

### Immunofluorescence

Sections were incubated with the following primary antibodies: anti-WT1 (Abcam, ab15249), anti-CD34 (Abcam, ab187282), anti-CD31 (Abcam, ab7388), anti-Ki-67 (CST, 11882), anti-Six2 (Abcam, ab220826), anti-endomucin (Abcam, ab106100), anti-CXCR4 (Abcam, ab181020), anti-NCAM1 (Abcam, ab220360), anti-FLK1 (Abcam, ab10972), anti-VEGFA (Abcam, ab39250). The primary antibodies were diluted 1:100 in immunofluorescence antibody dilution buffer (CST, 12378S), incubated at 4°C overnight and detected with Alexa Fluor 488 or 647 (Abcam) secondary antibodies. The secondary antibodies were diluted 1:600 in Immunofluorescence Antibody Dilution Buffer and incubated for 1.5 hours in a horizontal shaker at room temperature. Negative stainning control for each antibody was performed on kidney sections. Fluorescent images were photographed on Zeiss LSM780 confocal microscope.

## Abbreviations

FLK1: kinase insert domain receptor CD34
NCAM1: neural cell adhesion molecule 1
VEGFA: Vascular endothelial growth factor A
CXCR4: C-X-C motif chemokine receptor 4

## Declarations

## Acknowledgments

We thank Professor Bi-Sen Ding, Sichuan University, for providing CXCR4 Flox line.

## Funding

This work was funded by The National Key Research and Development Program of China (2017YFA0106600), the National Natural Science Foundation of China (No. 31771609, 81400747, 81270290).

## Ethics approval and consent to participate

All animal work has been conducted according to the ethics guidelines of the Army Medical University of China and and the experimental procedures were carried out in accordance with the Guide for the Care and Use of Laboratory Animals published by the National Institutes of Health (NIH Publication No. 85–23, revised 1996). This study was approved by the research ethics committee of the Army Medical University of China.

## Consent for publication

Not applicable.

## Competing interests

The authors declare no competing or financial interests.

## Additional Files

Additional file **1: Supplemental Fig. S1** Glomerular capillaries are formed in the same pattern before and after birth. (**a-c**) The development of glomerular capillaries during *E 13*. Capillary bed formation (**a**), capillary bed expansion (**b**) and the maturation stage (**c**). (**d-f**) The development of glomerular capillaries during *E 16*. Capillary bed formation (**d**), capillary bed expansion (**e**) and the maturation stage (**f**). (**g-i**) The development of glomerular capillaries during the first day after birth. Capillary bed formation (**g**), capillary bed expansion (**h**) and the maturation stage (**i**). Scale bar = 10 μm.

Additional file 2: **Video S1.** Three-dimensional structure of the blood vessel sprouting around renal progenitor cell aggregates. Six2 staining (green) indicates the renal progenitor cell aggregate, and FLK1 staining (red) indicates vascular sprouting occurs in the fetal kidney (arrowheads). Scale bar = 10 μm.

Additional file 3: **Video S2.** Three-dimensional structure of multiple capillaries extended into the glomerular cleft. WT1 staining (green) indicates the podocytes, and FLK1 staining (red) indicates vascular sprouting (arrowheads). Scale bar = 10 μm.

Additional file 4: **Video S3.** Three-dimensional structure of a capillary bed enlarges its area by sprouting. WT1 staining (green) indicates the podocytes, and FLK1 staining (red) indicates vascular sprouting (arrowheads). Scale bar = 10 μm.

Additional file 5: **Video S4.** Three-dimensional structure of a capillary bed formed in S-shaped body. NCAM1 staining (green) indicates the S-shaped body, and FLK1 staining (red) indicates vascular sprouting (arrowheads). Scale bar = 10 μm.

Additional file 6: **Video S5.** Three-dimensional structure of a bowl shaped capillary bed. WT1 staining (green) indicates the podocytes, and FLK1 staining (red) indicates capillary bed. Scale bar = 10 μm.

Additional file 7: **Video S6.** Three-dimensional structure of a redundant blood vascular pruning from glomerular capillaries during maturation stage. WT1 staining (green) indicates the podocytes, and arrowhead indicates pruning vascular. Scale bar = 10 μm.

