## supplemental for "Analysis the Detailed Process of Glomerular Capillary Formation Using Immunofluorescence Perform With Ultrathick Section"

**Supplemental material**

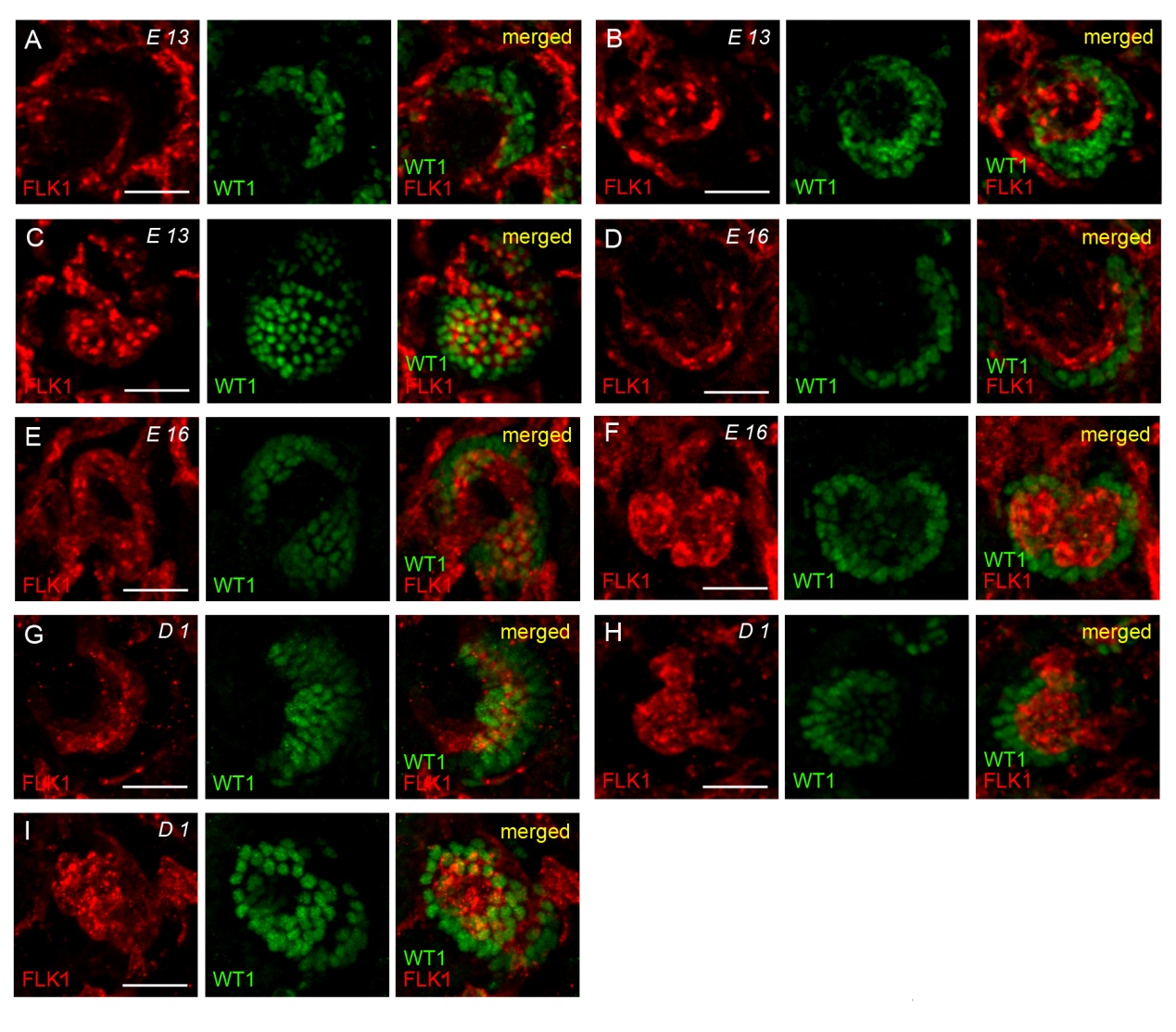

**Supplemental Fig. S1** Glomerular capillaries are formed in the same pattern before and after birth. (**a-c**) The development of glomerular capillaries during *E 13*. Capillary bed formation (**a**), capillary bed expansion (**b**) and the maturation stage (**c**). (**d-f**) The development of glomerular capillaries during *E 16*. Capillary bed formation (**d**), capillary bed expansion (**e**) and the maturation stage (**f**). (**g-i**) The development of glomerular capillaries during the first day after birth. Capillary bed formation (**g**), capillary bed expansion (**h**) and the maturation stage (**i**). Scale bar = 10 µm.

**Supplemental Video lengends**

**Supplemental Video 1.** Three-dimensional structure of the blood vessel sprouting around renal progenitor cell aggregates. Six2 staining (green) indicates the renal progenitor cell aggregate, and FLK1 staining (red) indicates vascular sprouting occurs in the fetal kidney (arrowheads). Scale bar = 10 µm.

**Supplemental Video 2.** Three-dimensional structure of multiple capillaries extended into the glomerular cleft. WT1 staining (green) indicates the podocytes, and FLK1 staining (red) indicates vascular sprouting (arrowheads). Scale bar = 10 µm.

**Supplemental Video 3.** Three-dimensional structure of a capillary bed enlarges its area by sprouting. WT1 staining (green) indicates the podocytes, and FLK1 staining (red) indicates vascular sprouting (arrowheads). Scale bar = 10 µm.

**Supplemental Video 4.** Three-dimensional structure of a capillary bed formed in S-shaped body. NCAM1 staining (green) indicates the S-shaped body, and FLK1 staining (red) indicates vascular sprouting (arrowheads). Scale bar = 10 µm.

**Supplemental Video 5.** Three-dimensional structure of a bowl shaped capillary bed. WT1 staining (green) indicates the podocytes, and FLK1 staining (red) indicates capillary bed. Scale bar = 10 µm.

**Supplemental Video 6.** Three-dimensional structure of a redundant blood vascular pruning from glomerular capillaries during maturation stage. WT1 staining (green) indicates the podocytes, and arrowhead indicates pruning vascular. Scale bar = 10 µm.
